# Evolution of affinity between p53 and MDM2 across the animal kingdom demonstrates high plasticity of motif-mediated interactions

**DOI:** 10.1101/2023.01.26.525693

**Authors:** Filip Mihalic, Emma Åberg, Pouria Farkhondehkish, Niels Theys, Eva Andersson, Per Jemth

## Abstract

The interaction between the transcription factor p53 and the ubiquitin ligase MDM2 results in degradation of p53 and is well studied in cancer biology and drug development. Available sequence data suggest that both p53 and MDM2-family proteins are present across the animal kingdom. However, the interacting regions are missing in some animal groups, and it is not clear whether MDM2 interacts with, and regulates p53 in all species. We used phylogenetic analyses and biophysical measurements to examine the evolution of affinity between the interacting protein regions: a conserved 12-residue intrinsically disordered binding motif in the p53 transactivation domain (TAD) and the folded SWIB domain of MDM2. The affinity varied significantly across the animal kingdom. The p53TAD/MDM2 interaction among jawed vertebrates displayed high affinity, in particular for chicken and human proteins (*K*_D_ around 0.1 μM). The affinity of the bay mussel p53TAD/MDM2 complex was lower (*K*_D_ = 15 μM) and those from a placozoan, an arthropod and a jawless vertebrate were very low or non-detectable (*K*_D_ > 100 μM). Binding experiments with reconstructed ancestral p53TAD/MDM2 variants suggested that a micromolar affinity interaction was present in the ancestral bilaterian animal and was later enhanced in tetrapods while lost in other linages. The different evolutionary trajectories of p53TAD/MDM2 affinity during speciation demonstrate high plasticity of motif-mediated interactions and the potential for rapid adaptation of p53 regulation during times of change. Neutral drift in unconstrained disordered regions may underlie the plasticity and explain the observed low sequence conservation in transactivation domains such as p53TAD.

**Statement for broader audience:** The protein p53 regulates central cellular processes including cell division and programmed cell death. p53 is regulated by another protein, MDM2, which binds to p53 and marks it for destruction. We measured the interaction between present-day and reconstructed ancient p53 and MDM2 proteins and found a range of binding strengths. Our findings suggest that rapid evolution of the p53/MDM2 interaction facilitates adaptation of p53 regulation during speciation.

## INTRODUCTION

The transcription factor p53 is involved in many cellular processes including cell cycle regulation and apoptosis ^1^. Because of this, mutations leading to p53 malfunction are highly correlated with cancer ^2^. The central position in cell cycle regulation is consistent with an ancient origin of p53, as evident from the presence of genes encoding p53-like proteins across the animal kingdom ^3–5^. It has been shown in well-studied vertebrate animals that p53 is kept at appropriate levels in healthy cells through ubiquitination and subsequent degradation ^6^. The ubiquitination process is initiated by binding of the intrinsically disordered p53 transactivation domain (TAD) to the SWIB domain of the ubiquitin ligase MDM2. The interaction is mediated by a conserved approximately 12-residue motif within p53TAD that folds into an α-helix upon binding to MDM2 (**Fig. 1A-B**).

**Figure 1.**
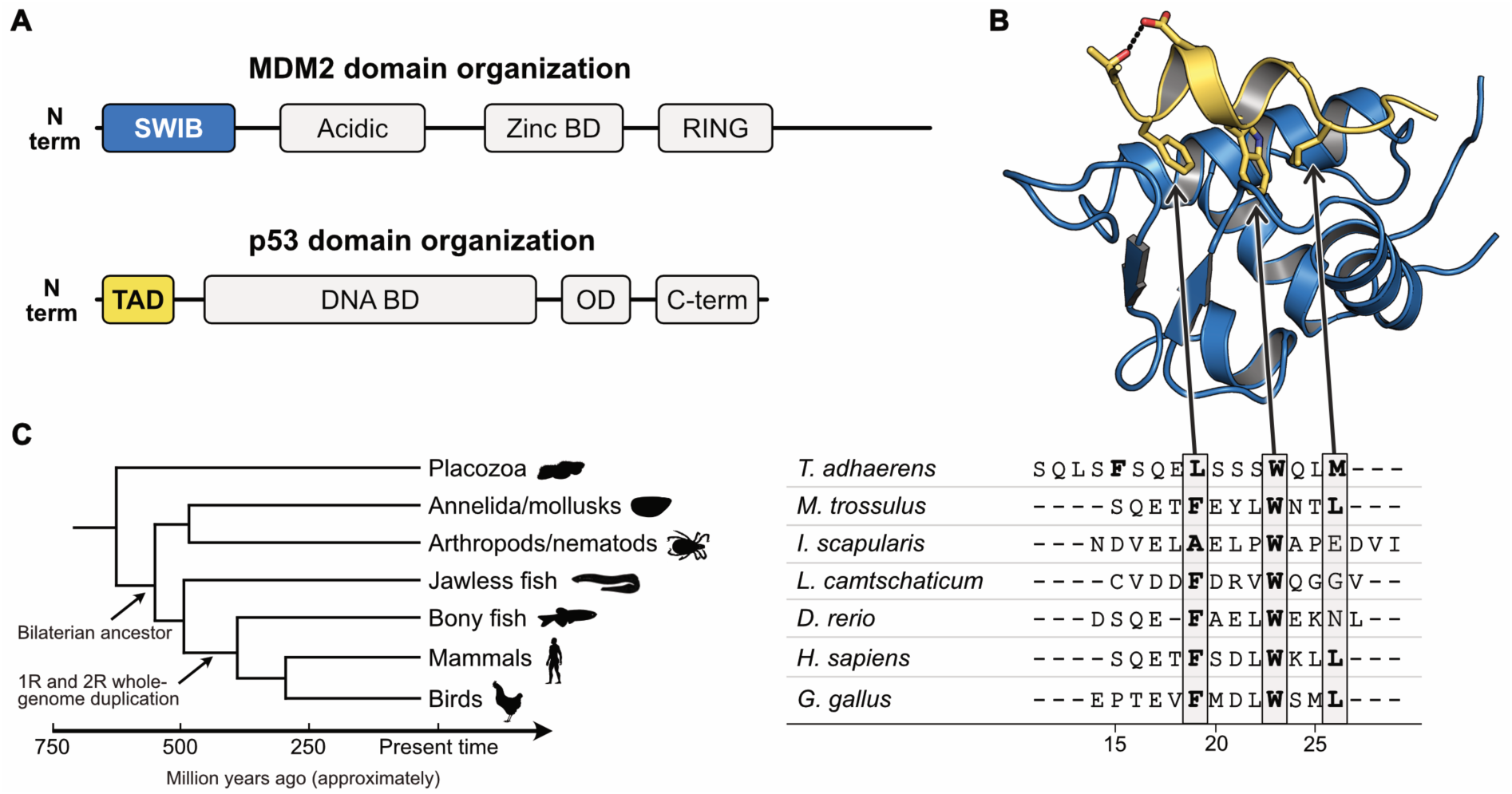
Sequences and structure of the p53TAD-MDM2 complex. **(A)** Schematic of p53 and MDM2 proteins with indicated folded domains. **(B)** Crystal structure of a complex between human p53TAD (yellow) and MDM2 (blue). The three conserved hydrophobic residues in the p53TAD motif, Phe19, Trp23 and Leu26 as well as Thr18 and Asp21 are shown as sticks. The hydrogen bond between Thr18 and Asp21 is shown as dots (PDB code: 1YCR) ^7^. **(C)** Schematic phylogenetic tree and sequence alignment of the most conserved region of p53TAD, which entails the canonical binding motif for MDM2, a 12-residue intrinsically disordered region that folds into an α helix upon binding to MDM2. The motif contains three hydrophobic residues that points straight down in the binding pocket of the MDM2 SWIB domain, namely F_19_xxxW_23_xxL_26_ in human p53TAD, highlighted in gray. The four residues N-terminal of the Phe residue, as well as Leu23 are also relatively well conserved. The numbering of motif residues is based on the human sequence throughout the paper to facilitate comparison. Note that an alternative alignment is possible for *T. adhaerens* p53TAD with a motif F_15_xxxL_19_xxxW_23_ that has one extra residue between Leu19 and Trp23. 1R and 2R represent two whole genome duplications in the vertebrate lineage. These genome duplications gave rise to the paralogs p53, p63 and p73 as well as MDM2 and MDM4 in extant vertebrates. The duplications might have occurred earlier than indicated, prior to or during ^8, 9^ the divergence of jawless fishes, but present day p53 paralogs in jawless fish appear not to correspond to those in other vertebrates. Animal silhouettes are from PhyloPic.

We have previously constructed phylogenetic trees of both the p53 and MDM2 protein families and examined their co-evolution across the animal kingdom with emphasis on their interaction domains p53TAD and SWIB ^10^. In agreement with earlier studies ^3, 11^ we found that the interaction domains are present in distantly related animal phyla, suggesting that the interaction between p53TAD and MDM2 dates back to the beginning of animal life. Furthermore, our analysis suggested that p53TAD and the SWIB domain of MDM2 is present in all extant deuterostome animals, but that both protein regions have disappeared in distinct protostome lineages, although splice variants ^12^ as well as coverage and quality of sequence databases complicate such analysis. In addition, while a SWIB domain-containing protein is present in *Drosophila*, it appears not to be part of an MDM2 homolog ^13^. Among the animals that diverged before the protostome/deuterostome split, the interaction domains were not found in cnidarians and porifera. However, they are present in the placozoan *Trichoplax adhaerens*, from an evolutionary branch that diverged before the radiation of all other extant animal phyla **(Fig. 1C)**^3^. Furthermore, pull down and co-transfection experiments suggested a functional p53/MDM2 interaction in *T. adhaerens* ^14^, in the protostome *Mytilus trossulus* (a mollusk) ^15^ and also in the non-jawed vertebrate Japanese lamprey ^16^. Thus, available data suggest an intriguing scenario where MDM2-dependent regulation of p53TAD was present in the last common ancestor of all animals and has been maintained in most extant animal phyla, but lost in some of them.

Prior to the radiation of extant vertebrate classes around 450 million years ago (Mya) two whole genome duplication events further shaped the evolution of the p53/MDM2 interaction. The genome duplications clearly occurred before the split of cartilaginous and bony fish ^17, 18^, but possibly even earlier, before or concurrent with the divergence of jawless fish ^8, 9^. In either case, the present-day paralogs p53, p63 and p73 as well as MDM2 and MDM4 likely originated in these genome duplications. The TAD in p53, p63 and p73 and the SWIB domain in MDM2 and MDM4 have been retained in extant vertebrate lineages, but evolution has subjected the proteins to sub- and neofunctionalization. Indeed, inspection of the amino acid sequences for vertebrate p53TAD orthologs reveals dramatic changes between different animal groups although a 12-residue binding motif is relatively well conserved (**Fig. 1C, Fig. S1**). Likewise, MDM2 and MDM4 display significant divergence with around 55 % amino acid identity for residues within the folded part of the SWIB domain, as defined by the crystal structure of human p53TAD/SWIB ^7^. A similar low identity is seen between human and lamprey MDM2 (53 %), consistent with a split of jawed (gnathostomes) and non-jawed vertebrates (agnatha, including extant lampreys) around the time of the whole genome duplications when the genes encoding MDM2 and MDM4 diverged ^19, 20^ (**Fig. S1**). In this paper we conform to the common naming of "p53" and "MDM2" also for non-vertebrate homologs, although it is not formally correct. In fact, the ancestral p53/p63/p73 protein was likely more similar to extant p63 and p73 than to p53 ^5^.

Conservation of sequence implies function. However, while phylogenetic methods are very powerful, affinity of protein interactions depends on fine molecular details and needs to be investigated by experiments. Therefore, since some animals apparently have lost their p53/MDM2 interaction while those that retained it display a significant divergence in amino acid sequence in the p53TAD interaction domains, we set out to directly assess the evolution of affinity between the conserved binding motif in p53TAD and the SWIB domain of MDM2 in the animal kingdom. Using biophysical measurements on extant as well as resurrected ancient proteins we found that the amino acid changes observed along different evolutionary trajectories have indeed modulated the affinity of the interaction. The history of the p53TAD/MDM2 interaction demonstrates the evolutionary plasticity and malleability of intrinsically disordered protein regions involved in protein-protein interactions.

## RESULTS

### p53TAD/MDM2 affinity across extant animals

Sequence-based predictions suggest that p53 and MDM2 were present in the last common ancestor of all animals ^3^. However, the interacting regions, p53TAD and the MDM2 SWIB domain could not be found in sequences from insects, crustaceans and cnidarians ^10^ or only in minor splice variants ^21^ suggesting that p53 is not regulated by MDM2 in these species. To further investigate the co-evolution between p53 and MDM2 we here mapped the interaction between p53 and MDM2 across the animal kingdom. Specifically, we measured affinity between the conserved 12-residue binding motif in p53TAD and the SWIB domain of MDM2 from extant animals representing different lineages that retained both interaction domains: mammals (human, *Homo sapiens*), birds (chicken, *Gallus gallus*), bony fishes (Zebra fish, *Danio rerio*), jawless fishes (Arctic lamprey, *Lethenteron camtschaticum*), arthropods (Deer tick, *Ixodes scapularis*), mollusks (Bay mussel, *Mytilus trossulus*) and the multicellular placozoan *Trichoplax adhaerens*. To facilitate comparison of residues at corresponding positions we use the numbering of human p53 residues based on a sequence alignment of the 12-residue binding motif (**Fig. 1C**). Sequences of all constructs used in the paper are compiled in **Supplementary Excel File 1**.

We used kinetics (stopped-flow spectrometry) or equilibrium experiments (fluorescence polarization, FP, or isothermal titration calorimetry, ITC) to determine the affinity of interactions between p53TAD and MDM2.

The affinity between p53TAD*_H.sapiens_* and MDM2*_H.sapiens_* was similar to that from previous studies (**Fig. 2A, Table 1**). Furthermore, human and chicken p53TAD/MDM2 complexes displayed similar affinity (80-100 nM) **(Fig 2B and D)**. In fact, p53TAD*_H.sapiens_*^15–26^ displays an apparently higher affinity (∼10 nM) with MDM2*_H.sapiens_*, but this is an artifact resulting from the short length of the peptide. Two longer variants, p53TAD*_H.sapiens_*^15–29^ and p53TAD*_H.sapiens_*^13–61^ have *K*_D_ values of 60-100 nM ^22^. This discrepancy has been previously reported by other groups ^23^ and suggested to be an effect of a non-native closing of the N-terminal lid in MDM2 over the binding cleft which is only possible with the shorter p53TAD peptide ^24^. Thus, while ∼100 nM is a good estimate of *K*_D_ for the native interaction between human p53TAD and MDM2, we use the short p53TAD*_H.sapiens_*^15–26^ (*K*_D_∼10 nM) for a direct comparison of binding to MDM2s from different species (**Table 2**). Note that several p53TADs display this "non-native" high affinity with MDM2*_H.sapiens_*.

**Figure 2.**
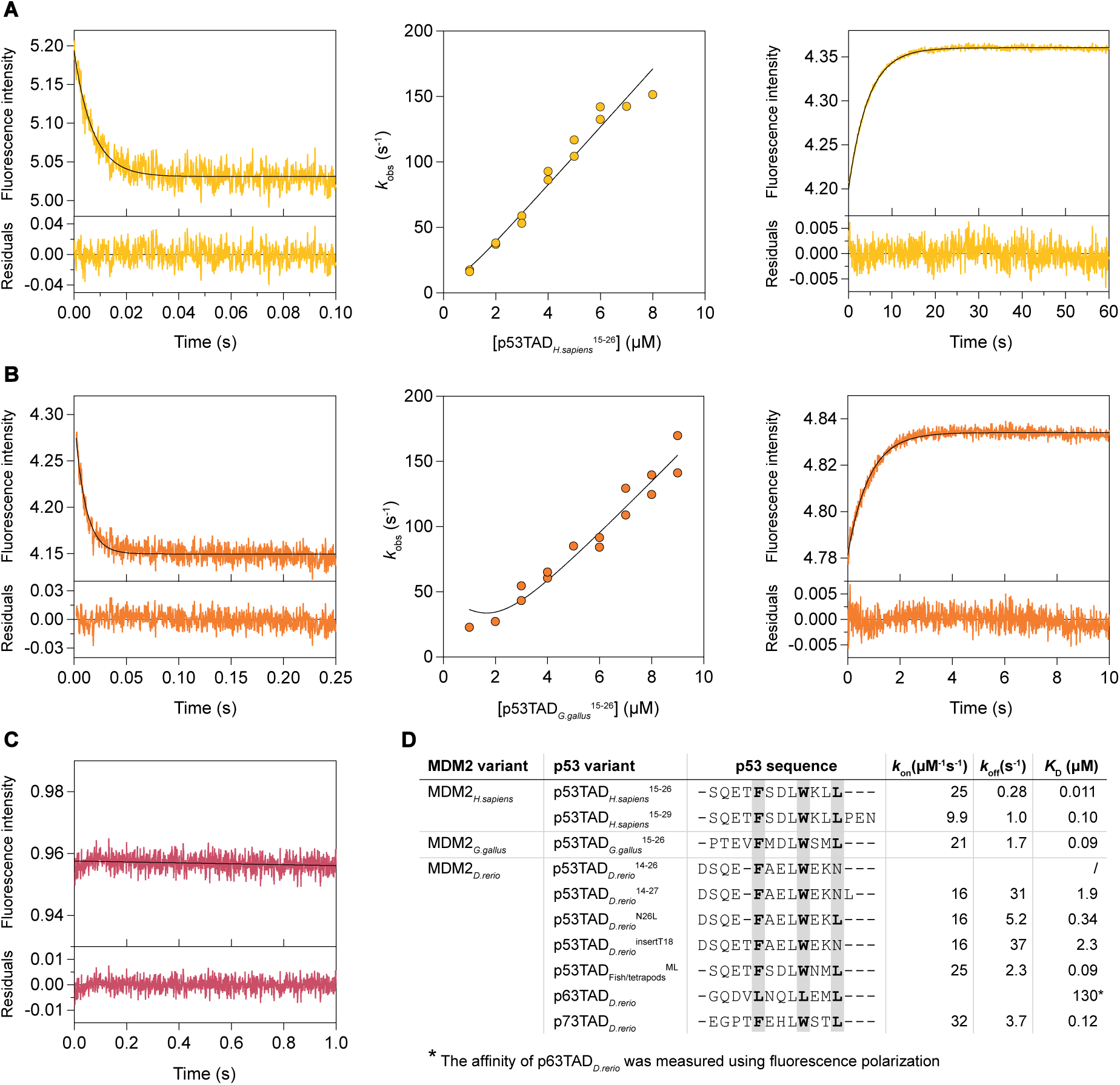
Affinity between p53TAD and MDM2 from extant vertebrate species. Binding was measured using stopped-flow spectroscopy. **(A)** Representative traces for the binding (left panel) and the displacement experiments (right panel) of MDM2*_H.sapiens_* interacting with p53TAD*_H.sapiens_*^15–26^. *k*_on_ is determined from the fitting of *k*_obs_ values obtained by performing association experiments at different peptide concentrations (middle panel). **(B)** Same experiments as in (**A)** for the chicken MDM2/p53TAD pair. **(C)** We could not observe any change in fluorescence upon mixing p53TAD*_D.rerio_*^14–26^ with MDM2*_D.rerio_*, indicating absence of interaction. **(D)** Sequence alignment and kinetic parameters for vertebrate MDM2-p53TAD interactions. The errors are reported in Table 1. *^a^*No binding could be detected with stopped flow spectroscopy. *^b^*The affinity of p63TAD*_D.rerio_* was measured using fluorescence polarization.

**Table 1.** Affinity and rate constants for resurrected and extant p53TAD-MDM2 interactions. The maximum likelihood (ML) contemporary (phylogenetic tree-matched) and extant interactions are highlighted in bold. AltAll represents alternative variants with all uncertain positions (posterior probability < 0.9) replaced with the second most likely amino acid residue. Point mutations are indicated as superscripts. Extant species in the table: zebra fish, *Danio rerio*; chicken, *Gallus gallus*; human, *Homo sapiens*; arctic lamprey, *Lethenteron camtschaticum*; bay mussel, *Mytilus trossulus*; deer tick, *Ixodes scapularis*); and the placozoa *Trichoplax adhaerens*. The p53TAD peptide corresponding to the conserved binding motif is 12 residues long and corresponds to human residues 15-26 unless otherwise specified.

| MDM2 variant | p53TAD variant | $K_D$ ($\mu\text{M}$) | $k_{\text{on}}$ ($\mu\text{M}^{-1}\text{s}^{-1}$) | $k_{\text{off}}$ ($\text{s}^{-1}$) |
| --- | --- | --- | --- | --- |
| **MDM2<sub>H.sapiens</sub>** | **p53TAD<sub>H.sapiens</sub><sup>15-26</sup>** | 0.011±0.002<sup>a</sup> | 25±1 | 0.28±0.03 |
| **MDM2<sub>H.sapiens</sub>** | **p53TAD<sub>H.sapiens</sub><sup>15-29</sup>** | 0.10±0.002<sup>b</sup> | 9.9±0.1 | 1.0±0.02 |
| **MDM2<sub>G.gallus</sub>** | **p53TAD<sub>G.gallus</sub><sup>14-26</sup>** | 0.09±0.01 | 21±1 | 1.7±0.1 |
| **MDM2<sub>D.rerio</sub>** | **p53TAD<sub>D.rerio</sub><sup>14-26</sup>** | -<sup>c</sup> |  |  |
|  | **p53TAD<sub>D.rerio</sub><sup>14-27</sup>** | 1.9±0.1 | 15.9±0.6 | 31±1<sup>d</sup> |
|  | p53TAD<sub>D.rerio</sub><sup>N26L</sup> | 0.34±0.01 | 15.5±0.3 | 5.2±0.1 |
|  | p53TAD<sub>D.rerio</sub><sup>insertT18</sup> | 2.3±0.4 | 16.1±0.6 | 37±5<sup>e</sup> |
|  | p53TAD<sub>Fishes/Tetrapods</sub><sup>ML</sup> | 0.093±0.01 | 24.6±0.8 | 2.3±0.2 |
|  | p73TAD<sub>D.rerio</sub> | 0.12±0.01 | 32±1 | 3.7±0.06 |
|  | p63TAD<sub>D.rerio</sub> | 130±2<sup>f</sup> |  |  |
| **MDM2<sub>L.camtschaticum</sub>** | **p53TAD<sub>L.camtschaticum</sub>** | 250±32<sup>f</sup> |  |  |
| **MDM2<sub>M.trossulus</sub>** | **p53TAD<sub>M.trossulus</sub>** | 15±3<sup>f</sup> |  |  |
| **MDM2<sub>I.scapularis</sub>** | **p53TAD<sub>I.scapularis</sub>** | -<sup>g</sup> |  |  |
| **MDM2<sub>T.adhaerens</sub>** | **p53TAD<sub>T.adhaerens</sub>** | -<sup>g</sup> |  |  |
| **MDM2<sub>Fishes/Tetrapods</sub><sup>ML</sup>** | **p53TAD<sub>Fishes/Tetrapods</sub><sup>ML</sup>** | 0.145±0.007 | 28±1 | 4.06±0.03 |
| MDM2<sub>Fishes/Tetrapods</sub><sup>AltAll</sup> | p53TAD<sub>Fishes/Tetrapods</sub><sup>AltAll</sup> | 0.098±0.005 | 30±1 | 2.93±0.07 |
| **MDM2<sub>Reptiles/Mammals</sub><sup>ML</sup>** | **p53TAD<sub>Reptiles/Mammals</sub><sup>ML</sup>** | 0.120±0.007 | 25.4±0.9 | 3.1±0.1 |
| MDM2<sub>Reptiles/Mammals</sub><sup>AltAll</sup> | p53TAD<sub>Reptiles/Mammals</sub><sup>AltAll</sup> | 0.20±0.01 | 18.6±0.6 | 3.7±0.2 |
| **MDM2<sub>Fishes/Tetrapods</sub><sup>ML</sup>** | **p53TAD<sub>1R</sub><sup>ML</sup>** | 0.66±0.08 | 14.3±0.4 | 9 ±1 |
|  | p53TAD<sub>1R</sub><sup>AltAll</sup> | 0.37±0.02 | 18.3±0.6 | 6.8±0.2 |
|  | p53TAD<sub>Bilateria</sub><sup>ML</sup> | 2.1±0.4 | 8.7±0.6 | 18±3 |
|  | p53TAD<sub>Bilateria</sub><sup>AltAll</sup> | 0.65±0.03 | 11.6±0.3 | 7.5±0.2 |

<sup>a</sup>The affinity for 12-mer p53TAD<sub>Human</sub> is incorrectly high, likely due to a non-native interaction. <sup>b</sup>Data acquired at $I = 0.20$ M from Åberg et al. <sup>22</sup>. <sup>c</sup>The affinity was too low to be measured by stopped flow spectroscopy. <sup>d</sup> $k_{\text{off}}$ from extrapolation = $28 \pm 4$ s<sup>-1</sup>. <sup>e</sup> $k_{\text{off}}$ from extrapolation = $32 \pm 3$ s<sup>-1</sup>. <sup>f</sup> $K_D$ from fluorescence polarization displacement experiment with low constant FITC-labeled p53TAD<sub>Human</sub>, which was displaced by excess p53TAD from the respective species. <sup>g</sup>Increase in fluorescence polarization signal was observed with FITC-labeled p53TAD<sub>Human</sub><sup>15-26</sup> (see Table 2), but the signal could not be reversed by unlabeled native p53TAD peptide suggesting non-specific binding of p53TAD<sub>Human</sub><sup>15-26</sup>. See Materials section for description of the errors.

**Table 2.**
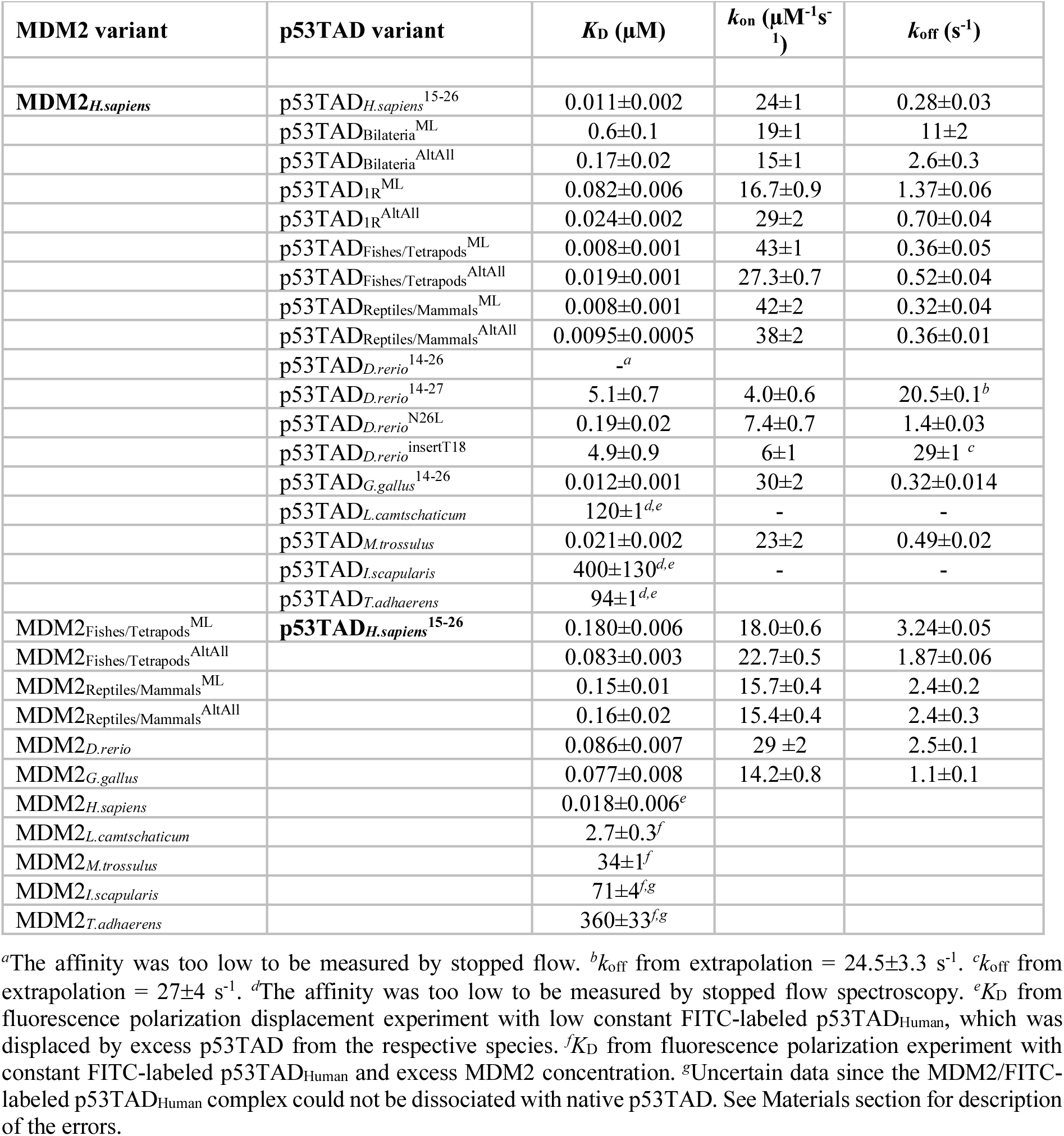
Affinity and rate constants for binding of human MDM2 to extant and ancient p53TAD and for binding of human p53TAD to extant and ancient MDM2 variants. The p53TAD peptide corresponding to the conserved binding motif is 12 residues long and corresponds to human residues 15-26 unless otherwise specified. The non-native interaction leading to high affinity between p53TAD_Human_^15–26^ and MDM2_Human_ is likely present also for the other 12-mer peptides such that relative affinities can be compared between the 12-mer peptides corresponding to human numbering 15–26.

Human and chicken p53TAD share the same hydrophobic triad F_19_xxxW_23_xxL_26_ (human p53 numbering) within the 12-residue binding motif, and their p53TAD/MDM2 complexes had similar affinity. However, there were significant differences in affinity of p53TAD/MDM2 from other animals. p53TAD*_D.rerio_*^14–26^, a 12-mer peptide corresponding to the conserved binding-motif region but with a deletion of Thr (or Ser) at position 18, showed non-measurable affinity for MDM2*_D.rerio_* with stopped flow spectroscopy. There is an apparent insertion of an Asn residue at position 26 in the conserved motif. Either changing Asn26 to Leu (p53TAD*_D.rerio_*^N26L^, *K*_D_ = 0.34 μM) or extending the peptide to a 13-mer to include Leu27 (p53TAD*_D.rerio_*^14–27^, *K*_D_ = 1.9 μM) resulted in detectable binding. Lack of a Thr18 helix N-cap also contributes to a lower affinity of the p53TAD*_D.rerio_*/MDM2*_D.rerio_* complex as shown by a peptide where Thr18 was introduced (p53TAD*_D.rerio_*^insertT^^18^, *K*_D_ = 2.3 μM) (**Fig. 2**, **Table 1**).

p53TAD/MDM2 complexes from protostomes and the non-jawed vertebrate *L. camtschaticum* (arctic lamprey) displayed much lower affinities than the bony vertebrates. For MDM2 from *M. trossulus* (bay mussel) and *L. camtschaticum* we could observe binding with a FITC-labeled human p53TAD^15–26^ peptide in FP experiments (**Fig. 3A, Table 1**). This labeled peptide could then be displaced by the respective unlabeled native p53TAD*_M.trossulus_* (*K*_D_ = 15 μM) or p53TAD*_L.camtschaticum_* (*K*_D_ = 250 μM), yielding estimates of the native *K*_D_ values **(Fig. 3B)**. However, we could not confirm these interactions by ITC (**Fig. S2**). Furthermore, we could not detect any binding between p53TAD and MDM2 from *T. adhaerens*, the most distantly related of all known animals, nor from the arthropod *I. scapularis* (deer tick), with either FP or ITC (**Fig. 3**, **Fig. S2**). For a specific interaction to the p53TAD-binding groove in MDM2, the unlabeled peptide is expected to compete out the labeled one. However, the signal did not decrease upon addition of unlabeled native p53TAD in the case of *T. adhaerens* and *I. scapularis* so the interaction with FITC-labeled p53TAD*_H.sapiens_*^15–26^ was therefore deemed non-specific **(Fig. 3B).**

**Figure 3.**
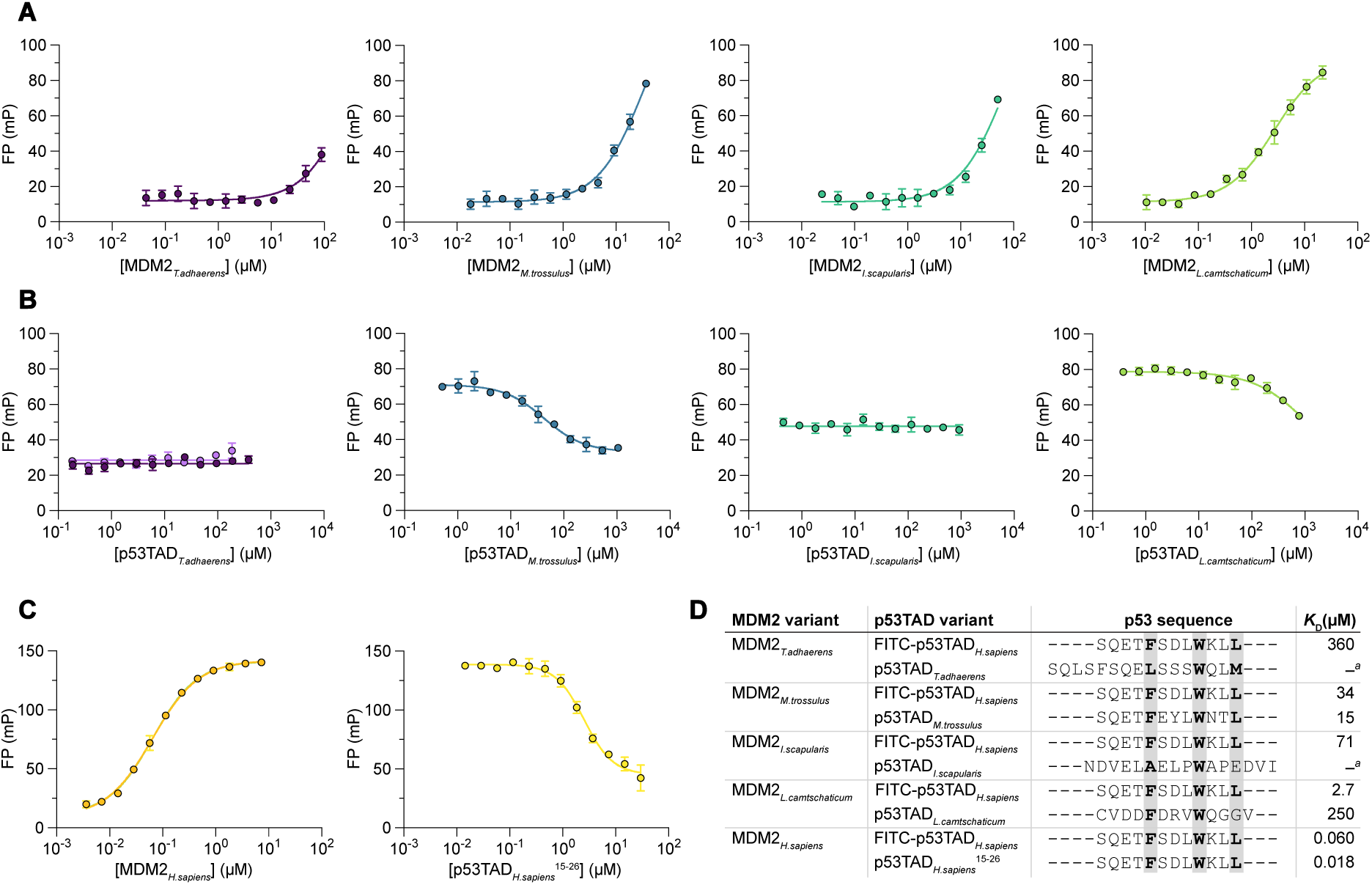
Affinity between p53TAD and MDM2 from non-vertebrate extant species. **(A)** Binding between MDM2s and FITC labeled p53TAD*_H.sapiens_*^15–26^ was measured using a fluorescence polarization-based saturation experiment. *K*_D_ for FITC labeled p53TAD*_H.sapiens_*^15–, 26^ was estimated from the curve fitting. **(B)** In the next step, FITC labeled p53TAD*_H.sapiens_*^15–26^ was competed out by an unlabeled peptide corresponding to the native p53TAD binding motif, as indicated on the x-axis. IC_50_ values from the resulting displacement data were used to estimate *K*_D_ of the native p53TAD/MDM2 interaction. For *T. adhaerens* displacement experiments, both a short p53TAD^11–26^ (dark purple) and a full length p53TAD (light purple, see **Supplementary Excel File 1**) were tested, but both failed to displace the weakly bound labeled peptide. **(C)** Saturation and displacement experiments showing the interaction between human MDM2 and p53TAD^15–26^ peptide for comparison. **(D)** Alignment of the binding motif of p53TAD from human, different non-vertebrate animal species and the jawless vertebrate arctic lamprey (*L. camtschaticum*). Approximate *K*_D_ values were determined from the data in panels **A**-**C**. Error bars for data points indicate standard deviation from technical replicates; n = 3. Errors for the parameters in panel **D** are shown in the **Supplementary excel file 2**. *^a^*No binding was detected.

A full-length version of p53TAD*_T.adhaerens_* (residues 1-126 of *T. adhaerens* p53) also did not show binding to MDM2*_T.adhaerens_* (**Fig. 3B**). We attempted expression and purification of full-length p53TADs from *I. scapularis*, *M. trossulus* and *L. camtschaticum*, but could not obtain enough of pure protein for binding experiments. One problem with absence of binding is that it is impossible to completely rule out misfolding of the MDM2 SWIB domain. To minimize the risk for misfolding, SWIB domains were subjected to circular dichroism (CD) and binding experiments immediately after purification. CD spectra and CD-monitored guanidinium chloride denaturation experiments suggested that all of the SWIB domains were soluble and folded (**Fig. S3**), yet they displayed weak or no binding to the corresponding native p53TADs.

Structural prediction of the complexes using ColabFold ^25^ suggested that p53TAD*_M.trossulus_*, p53TAD*_L.camtschaticum_* and p53TAD*_T.adhaerens_* all form an alpha helix upon binding to its respective native MDM2 (**Fig. S4**). p53TAD*_M.trossulus_* and p53TAD*_H.sapiens_* both have the same hydrophobic triad in the binding motif (FxxxWxxL) and the *M. trossulus* complex aligned well with the human complex. In agreement with the prediction, p53TAD*_M.trossulus_* bound MDM2*_H.sapiens_* with high affinity (**Table 2**). Close inspection of the MDM2*_M.trossulus_* ColabFold prediction revealed several key amino acid differences in the binding pocket as compared to MDM2*_H.sapiens_*, and offered an explanation for the low affinity of the native *M. trossulus* complex, despite the conserved canonical motif in p53TAD*_M.trossulus_*. These differences include Leu54 (Cys), Gly58 (Lys), Met62 (Gly) and Val93 (Phe) (numbering according to human MDM2 and with the *M. trossulus* residue in parenthesis; **Fig. S5**), all of which should interfere with optimal binding of the p53TAD. On the other hand, the low affinity of the *L. camtschaticum* complex is due to changes in the binding motif of p53TAD*_L.camtschaticum_* where only the first two hydrophobic residues align because of an extra residue between the conserved Trp and the third hydrophobic residue (FxxxWxxxV). p53TAD*_T.adhaerens_* displays two putative hydrophobic triads in the conserved region (FxxxLxxxW and LxxxWxxM). A predicted complex between a peptide containing the first motif and MDM2*_T.adhaerens_* aligned reasonably well with the human complex, while a peptide with the second motif was less well-defined with different directions of the peptide in the structural models (**Fig. S4**). Finally, p53TAD*_I.scapularis_* did not adopt the typical alpha helical conformation seen in the other complexes. Instead, p53TAD*_I.scapularis_* formed a more extended bound conformation where a Trp was found at the position of human Phe19, the conserved Leu aligned with human Trp23 and an Ala occupied the position corresponding to Leu26 resulting in a putative WxLxA motif with a reversed direction of binding compared to other p53TAD peptides (**Fig. S4C**). Importantly, the predictions generated for the *I. scapularis* and *T. adhaerens* complexes exhibited a markedly lower pIDDT confidence score, as defined by the Colabfold algorithm, with prediction certainty parameters of 50-60 compared to >90 for other complexes (**Fig. S4**). This low score suggested non-conservation of the binding motif consistent with the observed lack of affinity in the binding experiments.

### Phylogenetic reconstruction of ancient p53TAD and SWIB domain sequences

The large differences in affinity observed for the p53TAD/MDM2 interaction across the animal kingdom prompted us to investigate the evolution of affinity in greater detail in deuterostome lineages, where high affinity is observed (*K*_D_ = 100 nM-2 μM). The abundance of vertebrate sequences in public databases allowed for phylogenetic reconstruction of ancient sequences, which could then be expressed in *Escherichia coli*, purified and subjected to binding studies. Thus, to map the evolutionary trajectory of binding affinity between p53TAD and MDM2 we reconstructed and resurrected ancient contemporary (phylogenetic tree-matched) sequences in the jawed vertebrate lineage. Maximum likelihood (ML) ancestral protein sequences were predicted for the most recent common ancestor of different extant groups of organisms based on an alignment of proteins from extant species and their relationship in a phylogenetic tree. The amino acid positions with low probability in the reconstructed sequences (**Supplementary Excel File 3 and 4**) were analyzed with regard to their potential impact on binding using alternative variants denoted AltAll ^26^, in which all positions with a probability below a chosen threshold (0.90) were changed to the second most likely amino acid residue (if the probability was at least 0.10). The binding properties of this "unlikely but not impossible" variant was compared to those of the corresponding ML variant to assess whether the results were robust with regard to the uncertainty in the reconstruction.

We previously generated alignments and phylogenetic trees for both p53 and MDM2 family proteins ^10^ that were used in the current reconstruction. Following the whole genome duplications ≥450 Mya the vertebrate paralogs of MDM2 and MDM4 have obtained a substantial number of amino acid substitutions relative to each other resulting in a low sequence identity, which precluded reconstruction of the ancestral MDM2/MDM4 protein at 1R (**Fig. S1**). Thus, for MDM2, we reconstructed the sequence of the common ancestor of "reptiles" (including birds, turtles, crocodiles, snakes and lizards) and mammals living approximately 325 Mya, and for that of ray-finned fishes and tetrapods (∼420 Mya) (**Supplementary Excel File 3**).

Likewise, the p53TAD region has experienced substantial changes in terms of substitutions, insertions and deletions, even in evolutionarily recent times. In fact, the large number of changes precludes a reliable sequence alignment outside of the 12-residue region that facilitates binding to MDM2 SWIB (**Fig. 1, Fig. S1, Fig. S6, Fig. S7, Supplementary Excel File 4**). Thus, the 12-residue binding motif was reconstructed for the fishes/tetrapods and reptiles/mammals nodes, allowing a direct assessment of binding to phylogenetic tree-matched MDM2. We also reconstructed p53TAD ^ML^ and p53TAD ^ML^ from the time of the whole genome duplications 1R and 2R (∼450 Mya), corresponding to the ancestral p53/p63/p73 protein (1R) and p63/p73 (2R), respectively (and assuming that the duplications occurred in an ancestral gnathostome). Furthermore, we reconstructed the even older p53TAD_Bilateria_^ML^ from the common ancestor of all bilaterian animals (protostomes and deuterostomes) living ∼600 Mya.

### Evolution of the p53TAD/MDM2 affinity in the vertebrate lineage

Reconstructed ancestral protein sequences were resurrected through peptide synthesis or expression in *E. coli*. Peptides corresponding to p53TAD^ML^ and purified MDM2^ML^s and their AltAll versions were subjected to stopped-flow experiments in order to determine affinities. (**Fig. 4A, Table 1**). Finally, as a common reference, we measured affinities for ancestral p53TAD variants against MDM2*_H.sapiens_* and ancestral MDM2s against p53TAD*_H.sapiens_*^15–26^ (**Table 2**).

**Figure 4.**
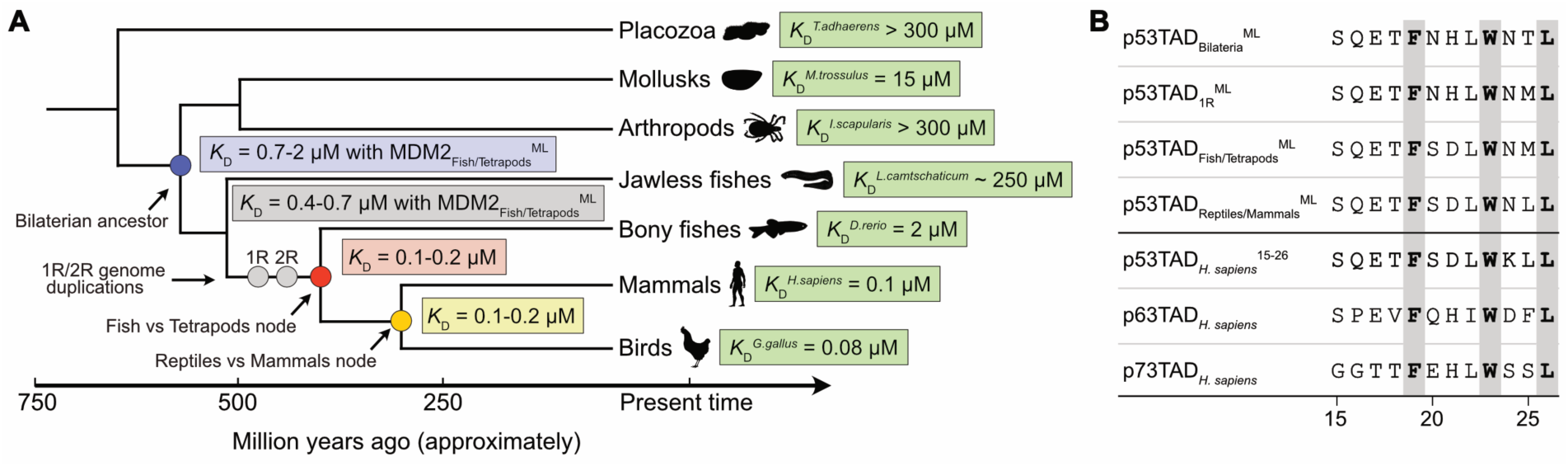
Evolution of MDM2-p53 interaction. **(A)** Simplified phylogenetic tree depicting the evolution of p53TAD/MDM2 affinity. *K*_D_ values for ancestral and extant complexes in selected lineages were determined by stopped flow spectroscopy and fluorescence polarization experiments (see Table 1 for errors). The p53TAD in the 1R node corresponds to the most recent common ancestral p53/p63/p73 protein reconstructed from extant jawed vertebrates (gnathostomes). The genome duplications giving rise to the p53 family paralogs may have occurred prior to the split with jawless fish (agnatha), but this putative earlier p53TAD_1R_ motif could not be reconstructed due to few agnatha sequences and unclear phylogeny of the *L. camtschaticum* paralogs. **(B)** Alignment of the reconstructed p53TAD binding motifs and the human p53TAD, p63TAD and p73TAD paralogs.

The binding affinity of the most ancient contemporary complex we could resurrect, that between MDM2_Fishes/Tetrapods_^ML^ and p53TAD_Fishes/Tetrapods_^ML^, was 145 nM (98 nM, for the AltAll complex) (**Table 1**). For MDM2_Reptiles/Mammals_^ML^ and p53TAD_Reptiles/Mammals_^ML^, we obtained a *K*_D_ of 120 nM (200 nM for the AltAll complex), which is very similar to that of the older fishes/tetrapod complex. Further diversification in the reptile and mammalian lineages retained or even increased the affinity of the complex, as shown by the native interactions of human and chicken p53TAD/MDM2 (**Table 1**), but also by the very similar (and artificially low) *K*_D_ values between MDM2*_H.sapiens_* and extant as well as ancient reconstructed p53TADs: p53TAD*H.sapiens*^15–26^, p53TAD*G.gallus*^14–26^, p53TADFishes/TetrapodsML, and p53TADReptiles/MammalsML (7.5-11 nM, **Table 2**). Furthermore, the ancestral MDM2_Fishes/Tetrapods_ and MDM2_Reptiles/Mammals_ display similar affinity with human p53TAD (*K*_D_ = 80-180 nM) as with their reconstructed contemporary p53TADs corroborating an overall retained affinity among tetrapods.

### Deeper evolution of the transactivation domain

The three paralogs in the gnathostome p53 family, p53, p63 and p73, probably originated from a gene present in an ancestral vertebrate at the time of the first whole genome duplication 1R ^19^. Generally, all three proteins have the three highly conserved hydrophobic residues F_19_xxxW_23_xxL_26_ within the 12-residue binding motif, although fish p63TADs usually contain Leu at position 23. However, outside of the binding motif, their TADs differ substantially in amino acid sequence following 450 Mya of evolution. In particular, p53TADs differ considerably between orthologs whereas p63TAD and p73TAD, respectively, are much more conserved (**Fig. S7**). Previous experiments demonstrated that p53TAD*_H.sapiens_* and p73TAD*_H.sapiens_* have high affinity for MDM2*_H.sapiens_* in contrast to p63TAD*_H.sapiens_*, which has low affinity (*K*_D_ = 6.1 μM) despite conservation of the F_19_xxxW_23_xxL_26_ motif ^22, 23, 27^. In the present study we included the p63 and p73 paralogs from *D. rerio*. Unlike p53TAD*_D.rerio_*, the conserved motif of its paralog p73TAD*_D.rerio_* retained high affinity towards MDM2*_D.rerio_* (*K*_D_ = 0.12 μM). p73TAD*_D.rerio_* has the F_19_xxxW_23_xxL_26_ motif whereas p63TAD*_D.rerio_* has a Leu residue in all three positions and very low if any affinity for MDM2*_D.rerio_* (*K*_D_ > 100 μM as estimated by fluorescence polarization) (**Figure 2D**). While there is some remaining redundancy in function, p53, p63 and p73 provide an example of gene duplication(s) followed by neofunctionalization, *i.e*., divergent evolution resulting in new functions where p53 is more involved in cell cycle arrest and apoptosis while p63 and p73 play a role in development ^28–30^.

We could not resurrect 1R and bilaterian MDM2 variants. Therefore, the affinities of the reconstructed p53TAD ^ML^ and p53TAD ^AltAll^ (from the most recent ancestral p53/p63/p73 protein) and of p53TAD_Bilateria_^ML^ and p53TAD_Bilateria_^AltAll^ (from the most recent ancestral bilaterian p53), were measured towards MDM2*_H.sapiens_* and MDM2_Fishes/Tetrapods_^ML^ (**Table 1, Table 2**). p53TAD_1R_ showed higher affinity for both MDM2*_H.sapiens_* (*K*_D_ = 24-82 nM) and MDM2_Fishes/Tetrapods_^ML^ (*K*_D_ = 370-660 nM) than p53TAD_Bilateria_ (*K*_D_ = 170-600 nM and 1.3-1.7 μM, respectively). Thus, in particular p53TAD_Bilateria_ binds much weaker than the historically younger p53TAD*H.sapiens*^15–26^, p53TAD*G.gallus*^14–26^, p53TADFishes/TetrapodsML, and p53TADReptiles/MammalsML. Comparison of sequences (**Fig. 4B, Supplementary Excel File 4**) suggests that two substitutions underlie the increased affinity of the younger p53TADs. First, Ser (or Thr) at position 25 was replaced by the hydrophobic residue Met (or possibly Gln), before the genome duplications and later with Leu in the tetrapod lineage. Second, the crystal structure of the human p53TAD/MDM2 interaction shows that Thr18 in p53TAD hydrogen bonds intramolecularly with Asp21, stabilizing the bound helix conformation ^7^. The present work suggests that His21→Asp was the original substitution that allowed formation of an N-cap, resulting in higher affinity, and which occurred between the genome duplications and the divergence of bony fishes and tetrapods. We also note that p53TAD ^ML^ (which is the most recent ancestral p63/p73 protein) was very similar to p53TAD ^ML^, but with a higher probability of Asp instead of Glu at position 17 (**Supplementary Excel File 4**), reflecting that the genome duplications 1R and 2R likely occurred relatively close in time, irrespective of when the agnatha lineage branched off.

## DISCUSSION

The evolution of functional traits is very complex and depends on interactions between multiple proteins. Despite the complexity, deciphering the molecular evolution of proteins can inform about structure-function relationships and evolutionary mechanisms governing their emergence ^31–33^. Intrinsically disordered regions ^34^ in proteins often contain motifs recognized by folded interaction domains ^35, 36^. The interaction between disordered binding motifs and domains are common in signaling pathways and transcriptional regulation and several aspects make such protein-protein interactions involving disordered protein regions particularly interesting from an evolutionary perspective. For example, except for certain key positions, such interactions appear robust to changes in the amino acid sequence increasing the possibility for permissive neutral substitutions, which may allow new functions to evolve upon further mutation and selection ^37^. In addition, interactions of this type are often hijacked by viral proteins ^38^, potentially imposing additional selection pressure. While we here attempted to pinpoint historical amino acid substitutions shaping the affinity between the p53TAD binding motif and MDM2, as done for other short motifs ^39^, conformational heterogeneity among the bound conformations of p53TAD is also likely contributing to the variation in affinity. Such structural plasticity, previously observed in evolution of protein-protein interactions ^37, 40^, is likely connected to general frustration of binding involving disordered regions, where several possible conformations compete and where none is perfectly optimized ^41^.

Our present study suggests an ongoing and dynamic evolution of affinity in the interaction between the intrinsically disordered p53TAD and the folded SWIB domain of MDM2. In particular, we observe a wide range of affinities for the p53TAD/MDM2 complex among extant bilaterian animals. This raises the question about the affinity of the ancestral bilaterian p53TAD/MDM2 complex present 600 Myr. Since reconstruction of an ancestral bilaterian MDM2 was not possible, we cannot be confident about the affinity of the ancestral complex. However, based on experimental data obtained with the reconstructed ancient p53TAD_Bilateria_^ML^, p53TAD_Bilateria_^AltAll^ and MDM2_Fishes/Tetrapods_^ML^, and with the extant MDM2_H.sapiens_ we speculate that the conserved binding motif in the bilaterian complex had a *K*_D_ value in the low μM range. Subsequently, higher affinity evolved in the lineage leading to fishes and tetrapods by formation of an N-cap in the p53TAD helix between position 18 and 21, and new hydrophobic interactions formed by the residue at position 25. After the split of the paralogs p53, p63 and p73 in an ancestral vertebrate, further evolution occurred in the motif in the respective paralog. For example, human and *D. rerio* p73TAD have high affinity for MDM2 and can likely form the N-cap, whereas p63TAD lacks hydrogen bond donor/acceptors at the corresponding positions, which results in a shorter helix and lower affinity ^42^. Moreover, the canonical binding motif in p53TAD*_D.rerio_* has evolved a lower affinity for MDM2 by loss of the hydrogen bond donor at position 18. In addition, an Asn residue at position 26 in the conserved interaction motif of p53TAD*_D.rerio_*, instead of Leu, further reduces affinity for MDM2. Nevertheless, the regulatory interplay between p53 and MDM2 is present: Previous studies have shown that DNA damage activates p53 in *D. rerio* ^43^ and that knockdown of MDM2 in embryos is lethal due to p53 activation and apoptosis ^44, 45^. Turning to non-vertebrates, the p53/MDM2 system appears functional in mollusks ^15^ and in *T. adhaerens* ^14^ despite the very low or even absent apparent affinity between their p53TAD and MDM2 SWIB domains. The low affinity can be explained by either a degenerate binding motif in p53TAD (*L. camtschaticum*), a suboptimal binding pocket in MDM2 (*M. trossulus*) or both (*I. scapularis* and *T. adhaerens*). The question arises: How can the observed radical differences in p53TAD/MDM2 affinity across bilaterian animals be compatible with a similar function? Obviously, while pivotal for binding to occur, affinity from interactions formed in the binding interface is not the sole determinant of a protein interaction. Concentrations of the interacting proteins could be regulated to compensate for suboptimal affinity. Affinity could also be modulated by disordered regions or even domains outside of the central binding interface ^46, 47^. However, for the human interaction, the conserved binding motif is sufficient for high-affinity MDM2 binding with little contribution from flanking regions, including the second less-conserved binding motif AD2, which binds CBP/p300 ^22, 23, 48–50^. In addition, we observed no binding with a full-length p53TAD from *T. adhaerens*. Thus, the lack of measurable p53TAD/MDM2 affinity in proteins from *T. adhaerens* and deer tick as well as the low affinity in Japanese lamprey and bay mussel p53TAD/MDM2 are intriguing. It is possible that p53 is not controlled by MDM2 in all species, and that the observed low affinity is a relic from an earlier functional interaction in the most recent bilaterian ancestor. Indeed, in the insect *Drosophila melanogaster* and the nematode *Caenorhabditis elegans*, which both lack a gene encoding MDM2, p53 is regulated by other proteins ^13, 51, 52^, showing that a loss of MDM2-mediated p53 regulation is biologically feasible. It is therefore not unreasonable that the functional interaction between p53 and MDM2 has been lost in other arthropod subphyla closely related to insects, such as chelicerata (ticks, spiders), although the interaction domains are still present in the proteins.

The dynamic evolution of p53TAD affinity, as illustrated by the changes at position 18 and 25, is also visible in its phosphorylation sites. Phosphorylation of Thr18 has been shown to reduce affinity for MDM2 while increasing affinity for the transcriptional coregulators CBP/p300 ^49, 50^. Furthermore, phosphorylation of Thr18 is contingent on prior phosphorylation of Ser15 ^53^. In agreement with these findings, both Ser15 and Thr18 (or Ser18) are highly conserved among vertebrates, but, surprisingly, both have been lost in the bird clade (**Fig. S6**). Another example involving phosphorylation is an apparently recent innovation in the primate lineage at Thr55, positioned in the second interaction motif AD2, which is much less conserved than the N-terminal motif. In human p53, AD2 was recently shown to regulate DNA binding via an intramolecular interaction with the adjacent DNA-binding domain, thereby promoting specific over non-specific DNA binding ^54, 55^. Phosphorylation of Thr55 further modulates this interaction by increasing the affinity of AD2 for the DNA-binding domain and thereby impeding binding cooperativity for the tetrameric p53 to DNA ^56^. Thr55 (or Ser55) is present in our closest relatives (monkeys, lemurs and tarsiers). However, p53s from mouse, rat (Asp55) and dog (Glu55) have a permanent negative charge in the corresponding position, suggesting a constitutively high affinity for the DNA-binding domain (**Fig. S6**).

Overall, the large differences in the p53TAD sequence observed among extant animals as well as the observed differences in affinity between the binding motif and the SWIB domain of MDM2 demonstrate high evolutionary plasticity in p53TAD and in its interaction with MDM2. By inference, the interactions between p53TAD and the transcriptional coregulators CBP/p300 may also be subject to changes over evolutionary time, as suggested by the striking changes at regulatory phosphorylation sites in p53TAD. In general, transcription factors regulating development and morphological phenotype are conserved ^57^. It is interesting to contrast the low conservation of p53TAD with the much higher conservation in p63TAD and p73TAD (**Fig. S7**), which (in particular p63) are proposed to be involved in development ^28^ and protection of DNA in the germ line, the likely ancestral function ^58^. Apparently, following the genome duplications 450 Myr, in the ancestral gnathostome lineage, p53 evolved towards a new and central role in monitoring DNA damage in somatic cells. On the face of it, the high evolutionary rate of changes in the amino acid sequence in p53TAD, evident from the comparison of different extant tetrapods (**Fig. S6**), is hard to reconcile with its interactions with both MDM2 and CBP/p300, and the centrality of p53 as a pleiotropic hub protein involved in multiple pathways ^58^. However, recent studies have shown that the exact sequence of transactivation domains are often not crucial for binding coactivators as long as key elements are present in the binding motif, in particular hydrophobic residues (Trp, Leu) separated by negatively charged Asp or Glu ^59–62^. It has also been shown that hub proteins such as p53 that connect different biological "modules" (intermodule hubs) are generally less constrained and evolve faster than hub proteins regulating a specific biological process (intramodule hubs) ^63, 64^. Furthermore, it is likely that p53 has slightly different roles related to life cycle and lifespan even among vertebrates. Thus, considering these points, evolution of p53TAD by relative fast neutral drift in unconstrained disordered regions ^65^ may underlie functional adaptation in different lineages and explain the observed low sequence conservation in p53TAD and variation in affinity for MDM2.

## Experimental procedures

### Ancestral sequence reconstruction

Sequence alignment of p53 was conducted in Guidance using the MAFFT algorithm with advanced options where the pairwise alignment option set to localpair and max-iterate option set to 1000 iterations. The full alignment contained 342 species (**Supplementary Text File 1 and 2**). The alignment was modified by removing gaps with a gap tolerance set to 95%. For p53, the alignment of bird sequences was manually curated such that they grouped with reptiles, which are closer relatives than mammals according to the established tree of life. Furthermore, echinoderm, hemichordate and lamprey sequences were removed since they did not group according to the tree of life. A phylogenetic tree was generated in MEGA X using maximum likelihood and the JTT substitution model. The tree was rooted against p53 from *T. adhaerens* and used to reconstruct ancestral sequences. The MDM2 alignment was performed with ClustalO and included sequences from 123 species (**Supplementary Text File 3**). A maximum likelihood tree was generated (JTT, MEGA X) and rooted against MDM2 from *Callorhinchus milli*, a chondrichthyes (cartilaginous fish). The alignment and phylogenetic tree were updated for MDM2 using additional fish and bird sequences as compared to the previous study ^10^. Thus, ancestral reconstruction was performed using MEGA X to obtain maximum likelihood sequences for p53TAD and the SWIB domain of MDM2, which are presented with posterior probabilities in supplementary information (**Supplementary Excel File 3 and 4**). AltAll variants of the reconstructed sequences included residues for which the posterior probability of the ML residue was lower than 0.90 *and* where the probability for the second most likely residue was at least 0.10.

The reconstructed sequences for p53TAD from the most recent common ancestor of fishes and tetrapods (p53TAD_Fishes/Tetrapods_^ML^) showed high confidence in many positions such as the three key residues defining the core motif: F_19_xxxW_23_xxL_26_. On the other hand, the residues at positions 20, 21 and 25 were less certain (**Supplementary Excel File 4)**. Two p53TADs were included in reconstructed ancient complexes (where both p53TAD and MDM2 were reconstructed): p53TAD_Fishes/Tetrapods_^ML^ and p53TAD_Reptiles/Mammals_^ML^. The second most probable residues were included in p53TAD_Fishes/Tetrapods_^AltAll^ at two positions (Ser18 and Leu25) and at one position (Met18) in p53TAD_Reptiles/Mammals_^AltAll^. The ML sequences from 1R and 2R differed at one position and since p53TAD_2R_ represents the ancestral p63/p73 motif, only p53TAD_1R_ was used in the experiments, since it represents the ancestral gnathostome p53/p63/p73 motif. Both p53TAD_1R_ and the older p53TAD_Bilateria_ were reconstructed with less confidence than the more recent p53TAD_Fishes/Tetrapods_ and p53TAD_Reptiles/Mammals_.

The reconstructed MDM2 variants used in the present study correspond to residues 17-125 of human MDM2, *i.e*., the ones visible in the crystal structure. Out of these 109 residues, 29 residues had a posterior probability below 0.90 in the MDM2_Fishes/Tetrapods_^ML^ variant and six residues a probability below 0.50. The majority of the uncertain residues are in either the N- or C-terminus. In the MDM2_Reptiles/Mammals_^ML^ variant, twelve residues had a posterior probability less than 0.90 of which only one was below 0.50 (the last one in the sequence). AltAll variants contained 23 (MDM2_Fishes/Tetrapods_^AltAll^) and twelve (MDM2_Reptiles/Mammals_^AltAll^) substitutions as compared to their respective MDM2^ML^ variant (**Supplementary Excel File 3**).

### ColabFold predictions

We used the ColabFold: AlphaFold2 using MMseqs2 ^25, 66, 67^ to predict complexes between the SWIB domain of MDM2 and peptides corresponding to the conserved binding motif in p53TAD, from different species. The structures of MDM2 were similar to that of the crystal structure of the human MDM2/p53TAD complex ^7^. p53TAD peptides were often, but not always predicted to bind in a similar fashion as in the human complex. The predictions were used to aid sequence alignment and interpretation of binding data as described in the Results section.

### Protein expression and purification

Extant and reconstructed cDNA encoding MDM2 variants were purchased from GenScript in either a pSY10 plasmid resulting in a construct with an N-terminal NusA domain followed by a TEV protease site, a His-tag, a PreScission protease site and the MDM2 variant: NusA-TEV-His_6_-PreScission-MDM2, or a pETM33 plasmid resulting in His_6_-GST-PreScission-MDM2 construct. Thus, the MDM2 SWIB domains shown in **Fig. 1** contains GPGS or GPMG at the N-terminus after PreScission digestion. A complete list of constructs is provided in **Supplementary Excel File 1**. The cDNA was transformed into *Escherichia coli* BL21 (DE3) pLys or *Escherichia coli* BL21 (DE3) cells (Invitrogen) using heat-shock. The cells were grown in LB medium at 37°C and overexpression of the fusion protein was induced with 1 mM isopropyl-β-D-thiogalactopyranoside when the optical density at 600 nm reached 0.7-0.8. The induced cultures were incubated overnight in a rotary shaker at 18°C. Cells were centrifuged and the pellet resuspended in a binding buffer (400 mM sodium chloride, 50 mM sodium phosphate, pH 7.8, 10% glycerol) followed by sonication to lyse the cells. Thereafter, cells were centrifuged at 4°C to remove cell debris and the supernatant was filtered and loaded onto a Nickel Sepharose Fast Flow column (GE Healthcare) in the case of pSY10 constructs. The fusion protein was eluted using binding buffer with 250 mM imidazole and then further purified using size-exclusion chromatography on a Hi load 16/60 Sephacryl S- 100 column (GE Healthcare) in the binding buffer with pH adjusted to 7.4. The fusion protein was then cleaved with PreScission (produced in house) protease overnight at 4°C followed by a second run on the size-exclusion chromatography column to remove the NusA protein. in the case of pETM33 constructs, after removing cell debris the supernatant was loaded onto Pierce™ Glutathione Agarose beads (Thermo Scientific), washed with wash buffer (50 mM Tris, 300 mM NaCl, pH 7.8) and eluted in GST elution buffer (50 mM Tris, 300 mM NaCl, 10 mM reduced glutathione, pH 7.8). Fusion protein was cleaved with PreScission protease overnight at 4°C and applied to Nickel Sepharose Fast Flow column to remove the tag. To remove any residual impurities final step of size-exclusion chromatography on a Hi load 16/60 Sephacryl S-100 column (GE Healthcare) in the binding buffer with pH adjusted to 7.4 was employed. Purity of all samples was checked with SDS-PAGE and MALDI-TOF mass spectrometry and pure samples were dialyzed against the experimental buffer (20 mM sodium phosphate, pH 7.4, 150 mM NaCl, 1 mM TCEP). Protein concentration was determined by measuring the absorbance at 280 nm and using extinction coefficients calculated from the amino acid sequence.

All p53TAD peptides were ordered as acetylated peptides (Ontores Biotechnology, GL Biochem Shanghai Ltd and GeneCust (France)) and dissolved in the experimental buffer (20 mM sodium phosphate, pH 7.4, 150 mM NaCl). The peptide identity was checked with MALDI-TOF mass spectroscopy and the concentration was determined by measuring Trp absorbance at 280 nm.

### Kinetic experiments

Affinity from kinetics was determined from measured values of association and dissociation rate constants (*K*_D_ = *k*_off_/*k*_on_), which can be determined with high precision and accuracy in stopped-flow spectroscopy experiments ^22^. All experiments were performed at 10°C in 20 mM sodium phosphate, pH 7.4, 150 mM NaCl with addition of 1 mM of the reducing agent TCEP (experimental buffer). Kinetic experiments were performed in order to determine the association and dissociation rate constants, *k*_on_ and *k*_off_, and the dissociation equilibrium constant, *K*_D_ (*i.e*., as the ratio of the rate constants, *k*_off_/*k*_on_). All experiments were performed on an upgraded SX-17 MV stopped-flow spectrometer (Applied Photophysics). The binding was monitored using the change in emission of Trp23 in p53TAD which binds in a hydrophobic pocket of MDM2. The excitation wavelength was set to 280 nm and emission was monitored at 330±25 nm using an optical interference (band-pass) filter. The rate constants, *k*_on_ and *k*_off_, were determined in separate experiments. In both cases the change in fluorescence was recorded with time and fitted to a single exponential equation to extract the observed rate constant *k*_obs_. To determine the association rate constant, MDM2 was held at a constant concentration of 0.5-2 μM (depending on the affinity of the complex and the change in fluorescence upon binding) and mixed with varying p53TAD concentrations in the range 1-10 μM. The observed rate constant, *k*_obs_, was plotted against the p53TAD concentration and fitted to a reversible bimolecular interaction ^68^, from which *k*_on_ was extracted. To determine the dissociation rate constant *k*_off_, a pre-formed complex of p53TAD and MDM2 (1 μM:1 μM) was mixed with excess (10-20 μM) of an N-terminally dansylated human p53TAD peptide (*D*-ETFSDLWKLLP), which displaced unlabeled p53TAD in the complex. The observed rate constant *k*_obs_ was plotted against the concentration of dansylated p53TAD and *k*_off_ was determined at high concentrations of displacer, where it equals *k*_obs_. For low affinity complexes, where *k*_off_ was >20 s^-1^, *k*_off_ could be independently determined from binding experiments and shown to be similar to those from displacement experiments. *k*_on_ values are given with the curve fitting error and errors in *k*_off_ are standard deviation from two or three replicates (displacement experiment) or fitting error (extrapolation in binding experiment). Errors for *K*_D_ values (= *k*_off_/*k*_on_) were calculated from propagating the errors of *k*_off_ and *k*_on_.

### Fluorescence polarization experiments

FP can detect weak affinities (*K*_D_ values in the high μM range), although the accuracy is lower than for the other methods. In FP experiments we observed direct binding using a FITC-labeled p53TAD peptide and used unlabeled peptides containing the binding motif to displace the labeled one. FP experiments were performed at room temperature using the same buffer as kinetic experiments. For saturation experiments the FITC labeled human peptide was held at a constant concentration of 6 or 10 nM and mixed with the 1:1 dilution series of different MDM2 proteins (highest concentration of proteins were 89, 37, 50, 22 and 7.3 μM for *T. adhaerens, M. trossulus, I. scapularis, L. camtschaticum* and *H. sapiens* MDM2, respectively) in black, non-binding surface, flat bottom 96-well plates (Corning Life Sciences). The polarization was measured at excitation/emission wavelengths of 485/535 nm on a SpectraMax iD5 plate reader. Saturation binding curves were fitted to a hyperbolic binding equation to obtain *K*_D_. For the displacement experiments the concentration of labeled human peptide and MDM2 protein (p53TAD*_H.sapiens_*^15–26^ peptide at 6 or 10 nM, MDM2 proteins at 50, 25, 32, 10 and 2 μM for *T. adhaerens, M. trossulus, I. scapularis, L. camtschaticum* and *H. sapeins* MDM2, respectively) were held constant and mixed with the 1:1 dilution series of the competing peptides (highest concentrations of peptides were 382, 187, 1060, 924, 775 and 29.5 μM for *T. adhaerens, T. adhaerens* full length*, M. trossulus, I. scapularis, and L. camtschaticum* p53TAD, respectively). The resulting displacement curve was fit to a sigmoidal dose response (variable slope) equation to obtain an IC_50_ value, which was in turn used to calculate *K*_D_ values for the competing peptides as described previously ^69^ (**Supplementary Excel File 2**). Errors were calculated as standard error of mean (SEM) of calculated *K*_D_ values for the displacer peptides resulting from independent fit of IC_50_ values for the replicate measurements.

### Isothermal titration calorimetry

ITC experiments were performed with MicroCal iTC200 (GE Healthcare) at 25°C. To minimize buffer mismatch the MDM2 proteins and p53TAD peptides were dialyzed against experimental buffer prior to experiments. The concentrations of MDM2 proteins in the cell was 59.4, 57.2, 62, and 44 μM for *T. adhaerens, M. trossulus, I. scapularis, and L. camtschaticum* MDM2 respectively and the concentration of p53TAD peptides in the syringe was 594, 587, 645, and 440 μM for *T. adhaerens, M. trossulus, scapularis, and L. camtschaticum* p53TAD, respectively. The data were analysed using the built-in software and the two-state binding model was assumed.

### Circular dichroism spectroscopy

To assess the secondary structure content of various MDM2 domains, CD was monitored between 200 and 250 nm with a 1 nm bandwidth, scanning speed 50 nm/min and data pitch 1 nm. Experiments were performed on a J-1500 spectrometer (JASCO) in experimental buffer at 25°C and at 20 µM MDM2 concentration. To monitor protein unfolding the proteins were mixed with increasing concentration of guanidinium chloride (GdnCl) up to a final concentration of 6 M and the CD signal was measured at 222 nm. The data (CD signal versus GdnCl concentration) were analyzed according to a two-state unfolding mechanism to obtain *m*_D-N_ value, and [GdnCl] midpoint of denaturation ^70^. A cooperative sigmoidal denaturation suggested that the MDM2 proteins were folded in the experimental buffer.

## Acknowledgments

This work was funded by the Swedish Research Council (2020-04395) and the Knut and Alice Wallenberg foundation (Evolution of new genes and proteins, 2015.0069) to PJ.

## Author contributions

Conceptualization, analysis, writing: FM, EÅ, and PJ. Methodology: FM, EÅ, PF, NT and EA.

## Supplementary Information

### Supplementary Figures

**Supplementary Fig. S1.**
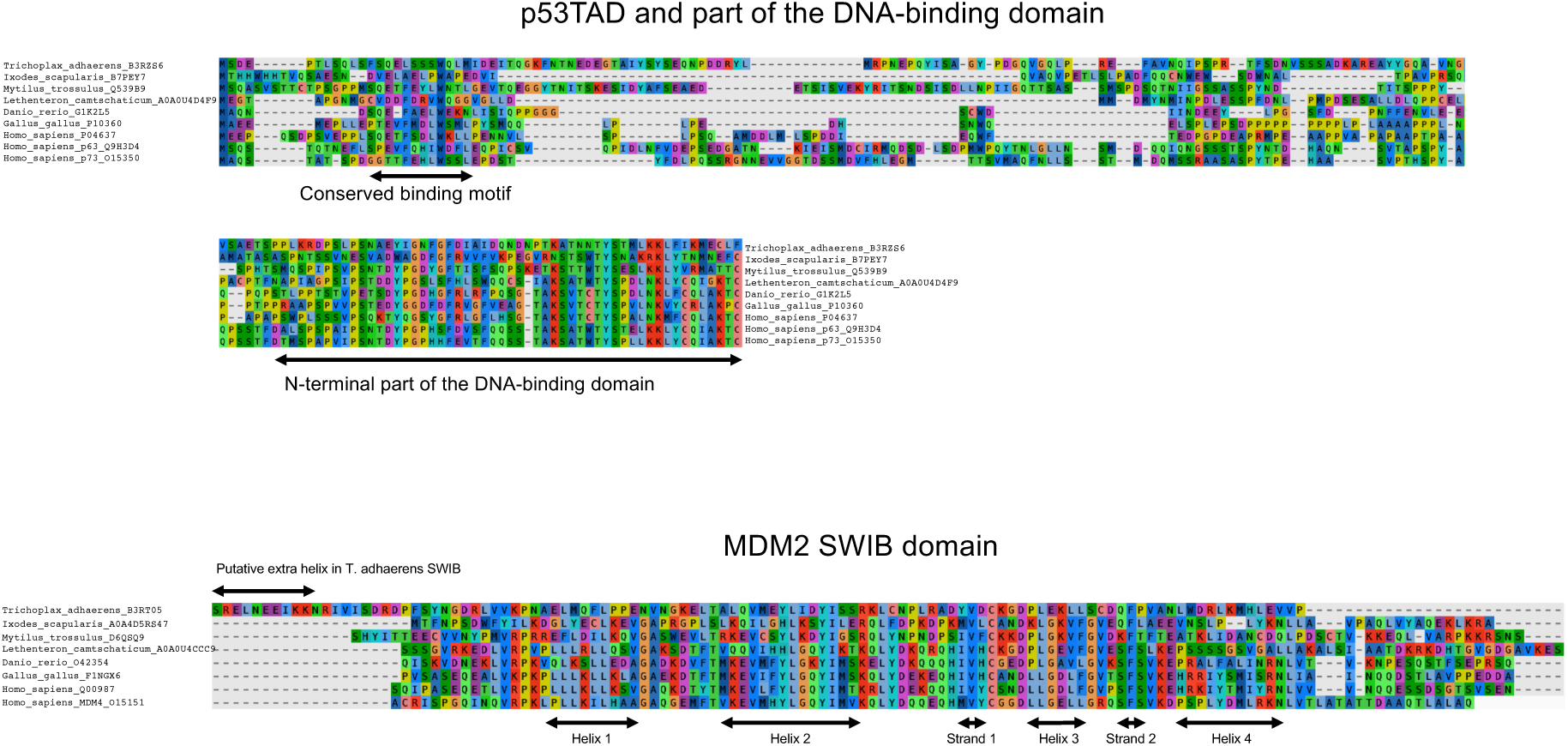
Sequence alignment of the p53 transactivation domain and MDM2 SWIB domain. **(A)** Sequence alignment of p53TADs from different species included in the study as well as the human paralogs p63 and p73. The sequence from the N-terminus to the beginning of the DNA-binding domain is shown for each protein. The alignment shows that the sequences have been subject to extensive evolution as reflected in the difference in amino acid sequences in proteins from extant animals. The large number of insertions and deletions precludes a correct alignment outside of the conserved binding motif. *Lethenteron camtschaticum* contains two additional p53 family proteins (Uniprot A0A0U4B546 and A0A0U3KDC1). The phylogeny of the three *L. camtschaticum* p53 paralogs in relation to gnathostome p53, p63 and p73 is not clear. Furthermore, the binding motifs appear to be lost in two of the paralogs and we have here aligned the sequence of the third one, with a remaining putative MDM2-binding motif. **(B)** Sequence alignment of the p53TAD-binding SWIB domains of MDM2 included in the study and of human MDM4. The sequence alignments were visualized with eBioX.

**Supplementary Figure S2.**
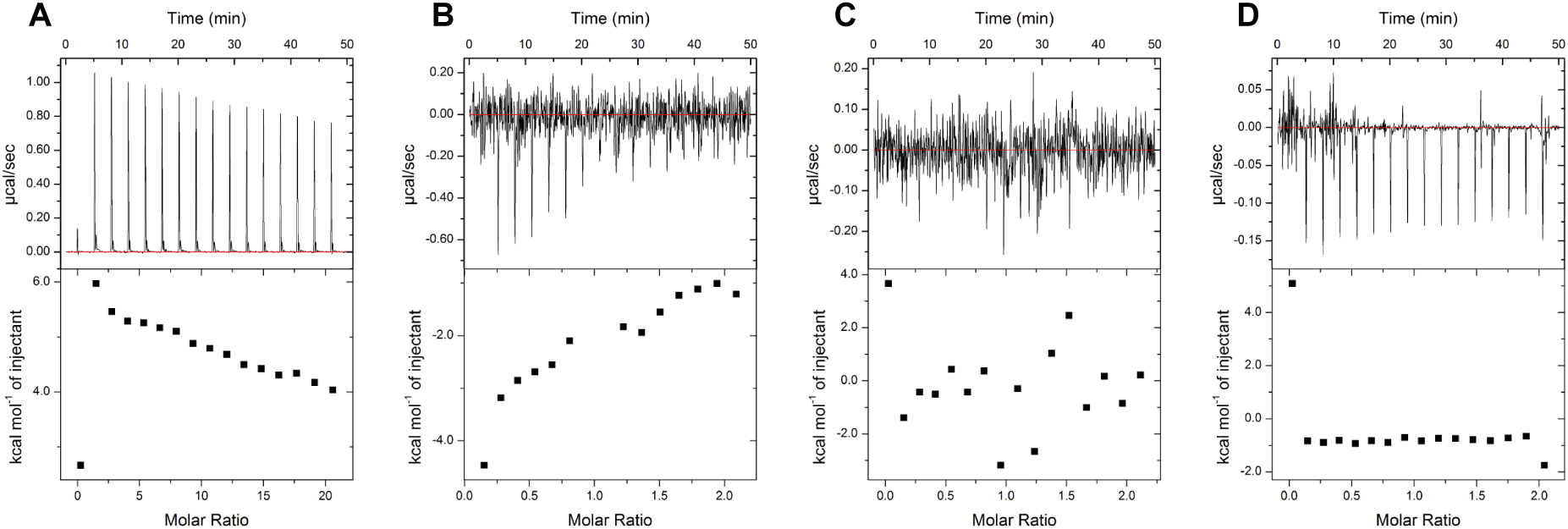
ITC experiments with p53TAD and MDM2 from different animals. Binding could not be detected between several p53TAD/MDM2 pairs. **(A)** p53TAD*_T.adhaerens_* was titrated into MDM2 *_T.adhaerens_*, **(B)** p53TAD*_M.trossulus_* was titrated into MDM2*_M.trossulus_*, **(C)** p53TAD*_I.scapularis_* was titrated into MDM2*_I.scapularis_* and **(D)** p53TAD*_L.camtschaticum_* was titrated into MDM2 *_L.camtschaticum_*.

**Supplementary Figure S3.**
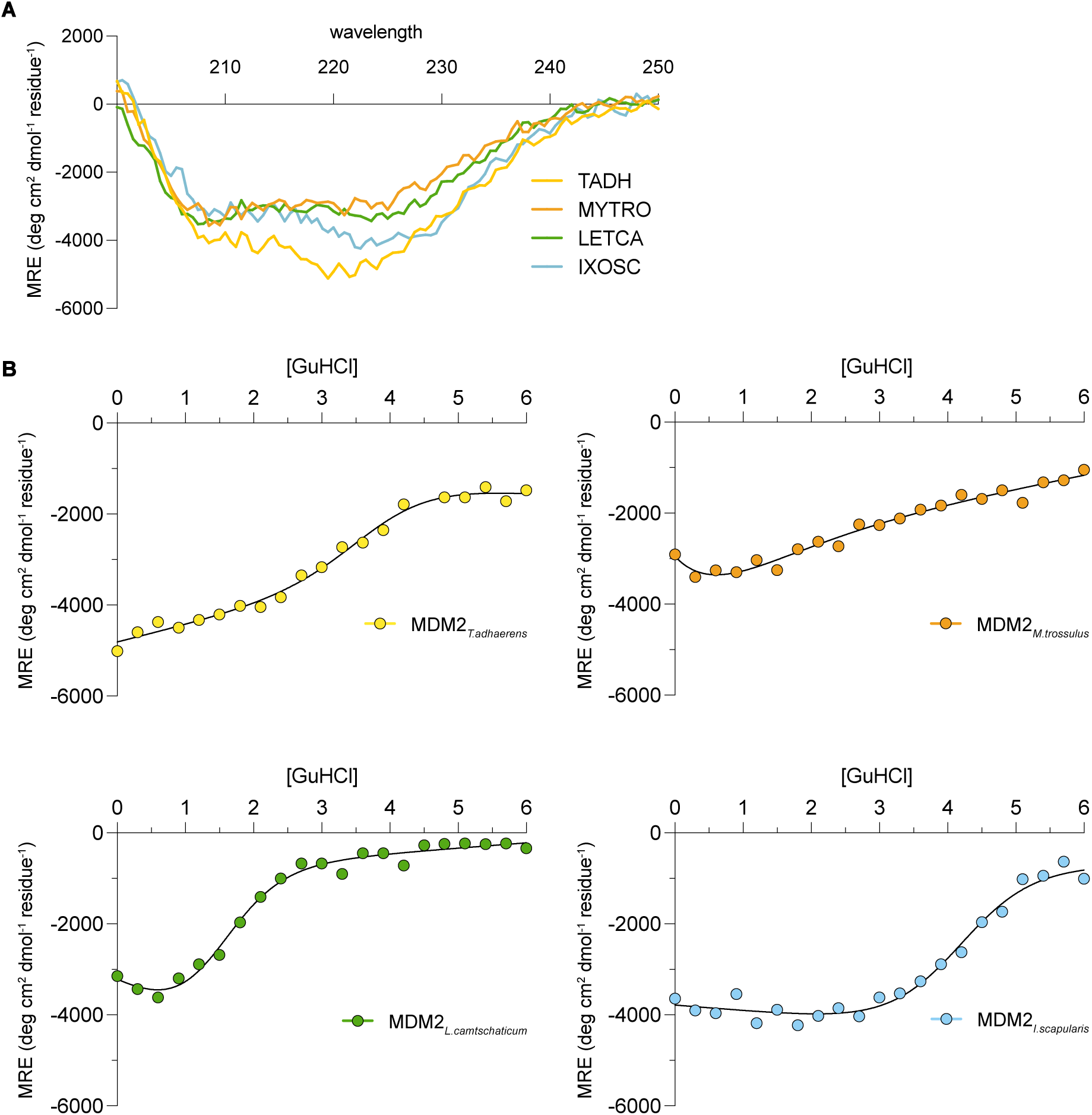
Thermodynamic stability of MDM2 variants. **(A)** Circular dichroism spectra of native and denatured MDM2 SWIB domain from *T. adhaerens*, *M. trossulus, L. camtschaticum* and *I. scapularis* respectively. **(B)** Guanidinium chloride-mediated denaturation of the domains monitored at 222 nm suggest that the proteins are folded. Solid lines correspond to fits to a two-state model. MRE, molar ellipticity.

**Supplementary Figure S4.**
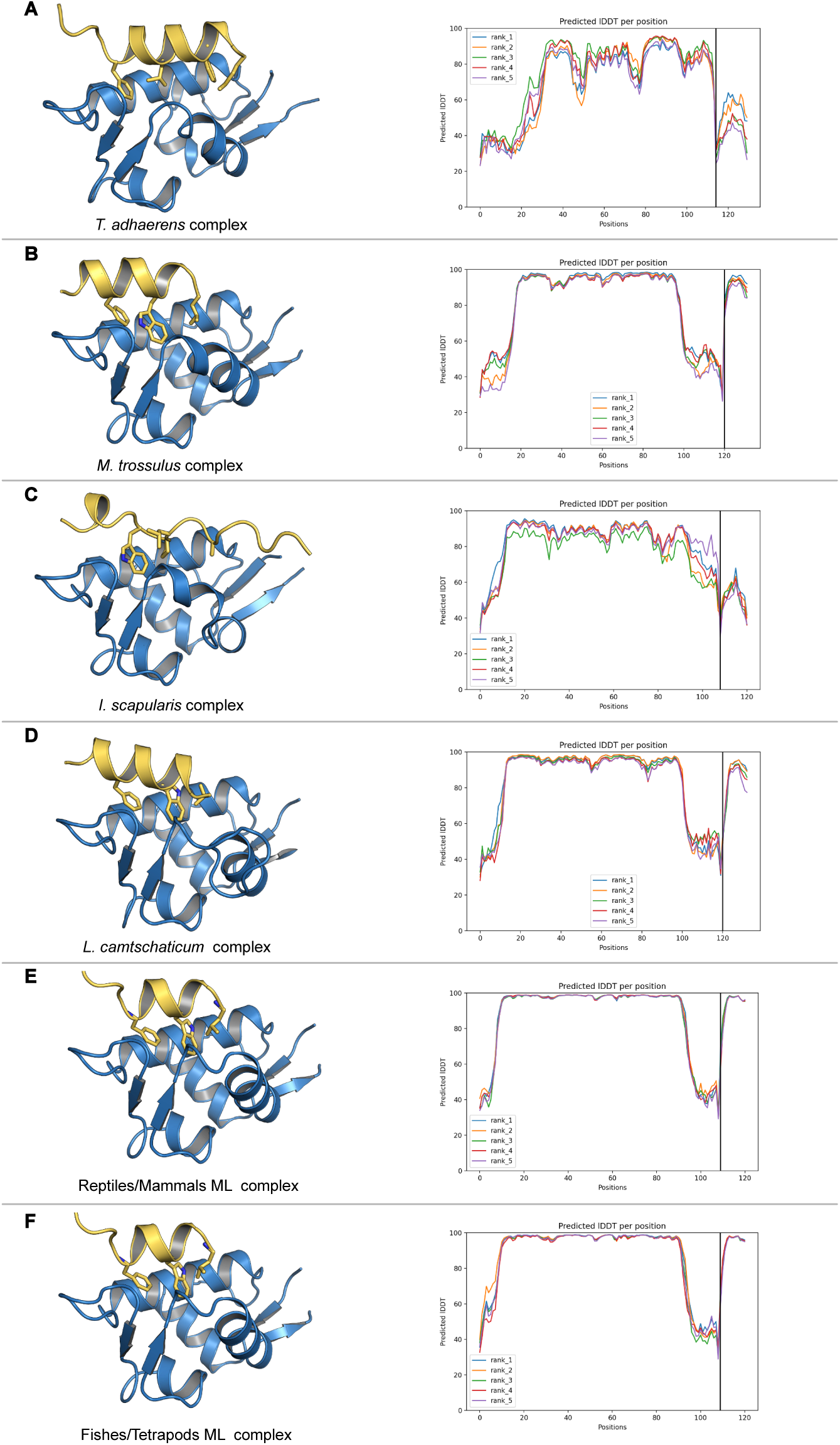
ColabFold predictions of p53TAD/MDM2 complexes. p53TAD is in yellow and MDM2 in blue. **(A)** Prediction for *T. adhaerens* complex with binding of the extended motif (FxxxLxxxWxxM). **(B)** Prediction for *M. trossulus* complex. **(C)** Prediction for *I. scapularis* complex. **(D)** Prediction for *L. camtschaticum* complex. **(E)** Prediction for the resurrected ancestral reptiles/mammals complex. **(F)** Prediction for the resurrected ancestral fishes/tetrapods complex. In all cases, the p53TAD residues Phe19, Trp23 and Leu26 pointing into the hydrophobic pocket are shown as sticks.

**Supplementary Figure S5.**
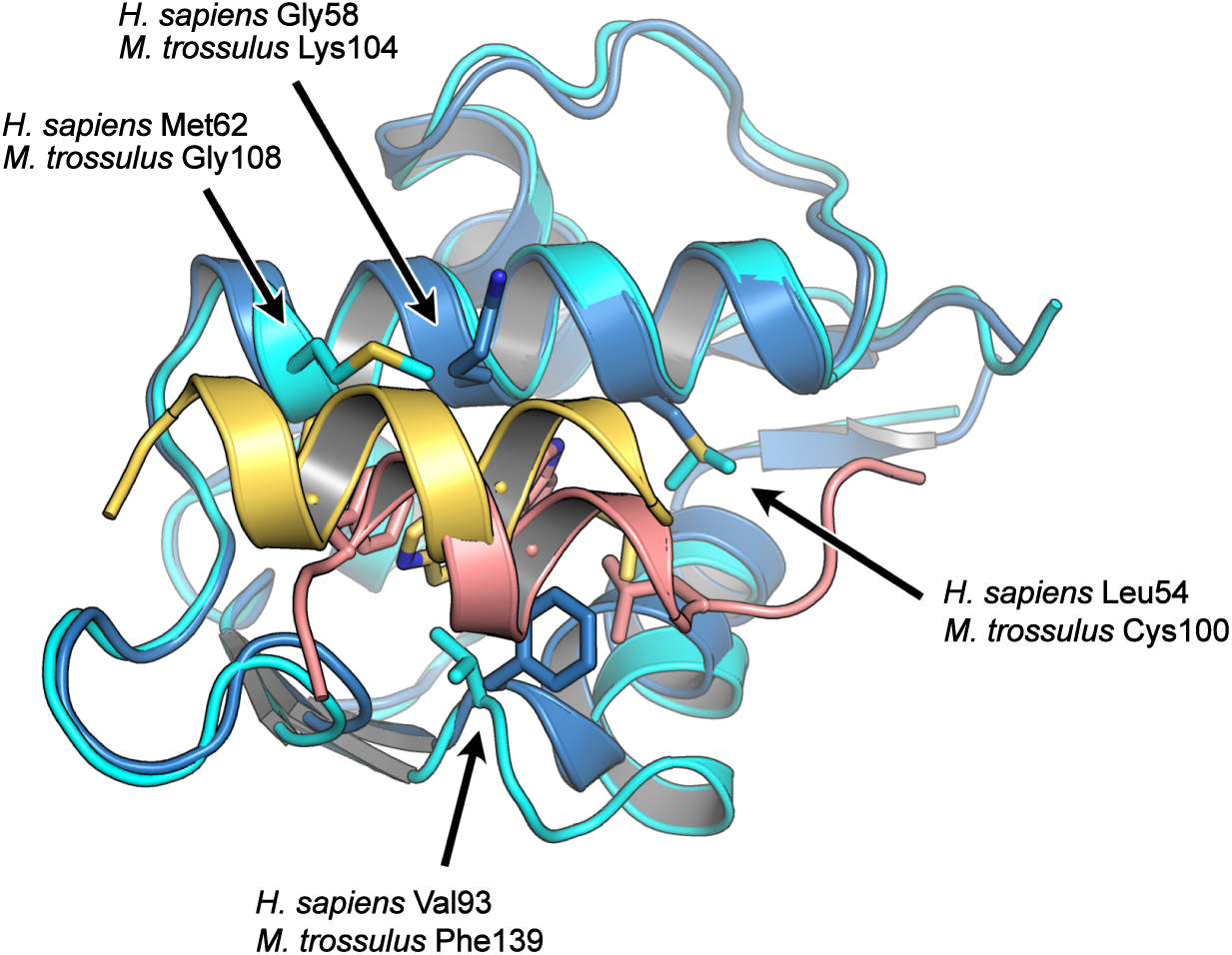
Colabfold prediction for the *M. trossulus* complex compared to the crystal structure model of the human complex. The human MDM2 is in cyan (PDB id: 1YCR) [7], human p53TAD in salmon, *M. trossulus* MDM2 in blue and *M. trossulus* p53TAD in yellow. Arrows indicate key differences between human and *M. trossulus* MDM2 that likely provide a hydrophobic p53TAD binding pocket with sub-optimal p53TAD binding properties (amino acid numbering according to human MDM2 protein).

**Supplementary Figure S6.**
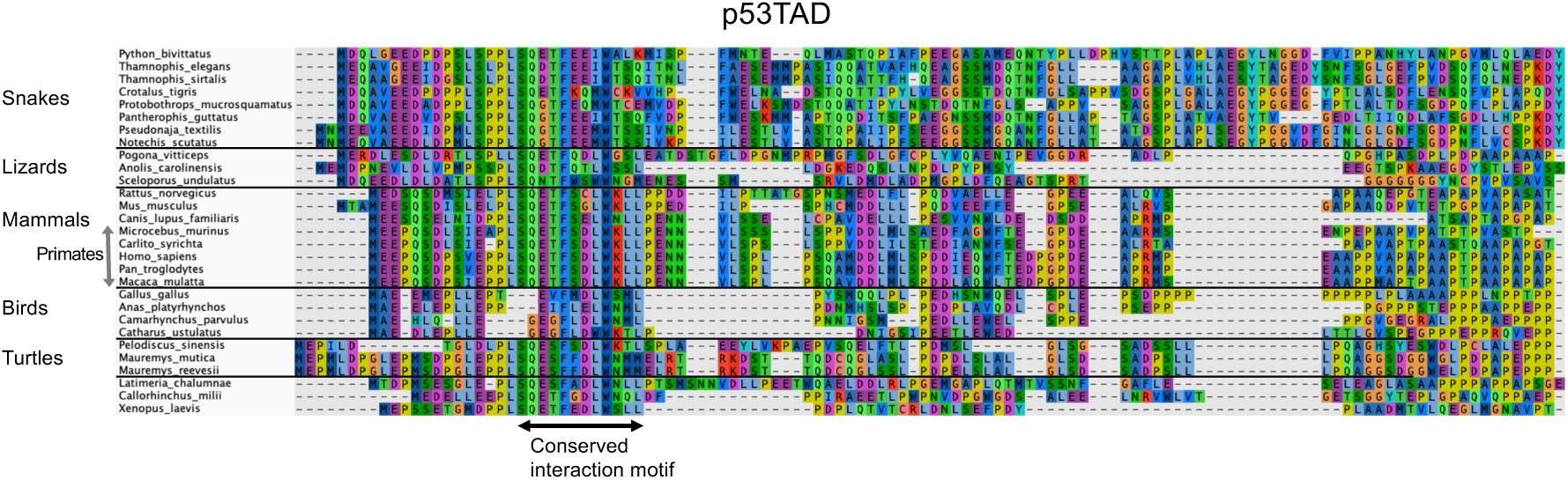
Sequence alignment of p53TAD from tetrapods. Alignment of p53TAD colored according to sequence similarity. p53TAD can only be confidently aligned for closely related species such as primates. The alignment was performed using Muscle and adjusted manually in the N-terminus for birds and *P. sinensis*. Note that both phosphorylation sites Ser15 and Thr18 have been lost in the bird p53s. The figure was made in AliView.

**Supplementary Figure S7.**
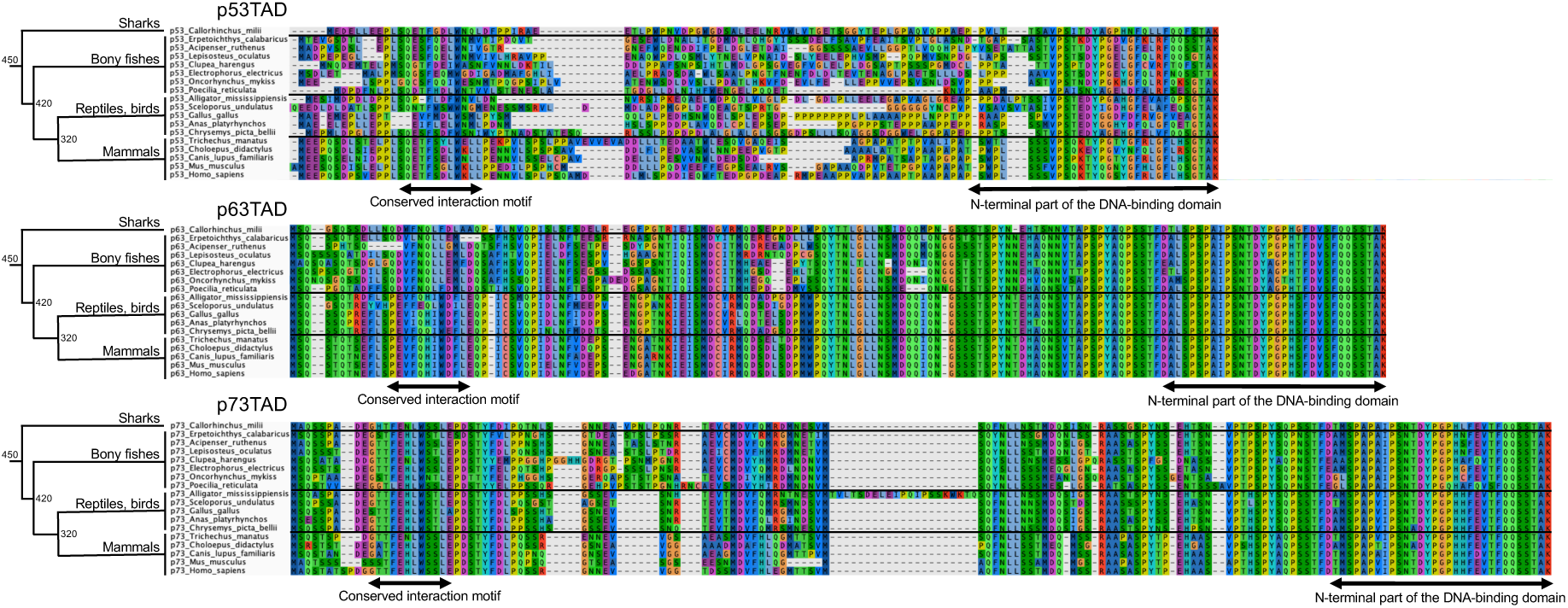
Sequence alignments of TADs from p53, p63 and p73 from vertebrates. It is clear that the TADs of p63 and p73 are more conserved than that from p53. For p63 and p73, the alignment starts at the Met residue closest to the canonical binding motif. Thus, annotations in Uniprot or NCBI may include additional residues in the N-terminus. The long insertion in *Alligator mississippiensis* p73TAD (TVLTSDELEIPQIPSSKWKTQ) is present in its crocodilian relative *Crocodylus porosus* but not in *Gavialis gangeticus* p73TAD. The alignment was performed using Muscle and the figure was made in AliView. The numbers at the nodes of the phylogenetic tree shows the approximate time of divergence in million years ago.

### Supplementary Tables, see separate files

**Supplementary Excel File 1. Sequences of constructs used in the experiments.** The sequence and name for each interaction motif in p53TAD and MDM2 SWIB domain used in binding experiments in the study.

**Supplementary Excel File 2. Calculation of *K*_i_ from IC50 values.** IC50 values were determined as described in the materials section. The theory behind the conversion of IC50 values to *K*_i_ values (=*K*_D_) is described in Nikolovska-Coleska, *et al.*, Development and optimization of a binding assay for the XIAP BIR3 domain using fluorescence polarization. *Anal. Biochem.* **332**, 261–273 (2004).

**Supplementary Excel File 3. Reconstruction of the SWIB domain of MDM2.** Reconstructed maximum likelihood (ML) and low probability AltAll versions of the MDM2 SWIB domain. The posterior probability is shown for each amino acid for each position for the reconstructed sequences.

**Supplementary Excel File 4. Reconstruction of p53TAD.** Reconstructed maximum likelihood (ML) and low probability AltAll versions of the conserved interaction motif in p53TAD. The posterior probability is shown for each amino acid for each position for the reconstructed sequences.

### Supplementary text files, see separate files

**Supplementary Text File 1. Fasta file for p53TAD.** The sequence alignment file used for ancestral reconstruction of the interaction motif in p53TAD.

**Supplementary Text File 2. Full length sequences of p53.** Full length sequences of p53s used in the reconstruction. This file contains sequences that were removed in the final reconstruction, such as echinoderms, hemichordates and agnatha (lampreys).

**Supplementary Text File 3. Fasta file for MDM2 SWIB.** The sequence alignment file used for ancestral reconstruction of the SWIB domain of MDM2.

